# Autophagy-cell death balance is maintained by Polycomb-mediated regulation during stem cell differentiation

**DOI:** 10.1101/2022.03.29.486206

**Authors:** Deepika Puri, Aparna Kelkar, Gauri Shankar Bhaskar, Deepa Subramanyam

## Abstract

Autophagy is a conserved cytoprotective process, aberrations in which, lead to numerous degenerative disorders. While the cytoplasmic components of autophagy have been extensively studied, the epigenetic regulation of autophagy genes, especially in stem cells, is less understood. Deciphering the epigenetic regulation of autophagy genes becomes increasingly relevant given the therapeutic benefits of small-molecule epigenetic inhibitors in novel treatment modalities. We observe that, during retinoic acid-mediated differentiation of mouse embryonic stem cells (mESCs), autophagy is induced, and identify the Polycomb enzyme EZH2 as a regulator of this process. In mESCs, EZH2 represses several autophagy genes including the autophagy regulator *Dram1*. EZH2 facilitates the formation of a bivalent chromatin domain at the *Dram1* promoter, which allows the expression of the gene and induction of autophagy during differentiation, while still retaining the repressive H3K27me3 mark. EZH2 inhibition leads to loss of the bivalent domain, and a consequential “hyper- expression” of *Dram1*, with extensive cell death. This study shows that Polycomb group proteins help maintain an autophagy-cell death balance during stem cell differentiation, in part, by regulating the expression of the *Dram1* gene.

## Introduction

Autophagy (macroautophagy) is an evolutionarily conserved catabolic process in which, cytosolic components are sequestered by double-membraned vesicles called autophagosomes, and transported to the lysosome for degradation [1, 2]. The mechanism of autophagy includes steps such as autophagy initiation, nucleation, autophagosome formation, fusion with the lysosome, and ultimately lysosomal degradation [1]. This involves the function of a distinct set of evolutionarily conserved genes called Autophagy-related genes (ATG) that form complexes with other proteins [3]. While being constitutively active at low levels, external stimuli such as starvation, stress, etc, lead to an induction of autophagy. Autophagy modulation facilitates cellular homeostasis, metabolic turnover, and degradation of organelles and misfolded proteins [4]. Defects in autophagy are associated with numerous degenerative disorders such as Alzheimer’s Disease, Parkinson’s Disease, Amyotrophic Lateral Sclerosis among others [5]. While autophagy was previously thought of as a process, purely regulated by changes in cytosolic components, recent studies highlight the role of transcriptional regulation of autophagy genes in governing this pathway [6, 7].

Epigenetic regulation, primarily mediated by modifications to chromatin, is one of the major governing factors of gene transcription [8]. Autophagy genes are regulated at the chromatin level by epigenetic modifiers such as BRD4 [9], G9A [10], CARM1 [11], and hMOF [12]. Additionally, transcription factors such as TFEB, FOXO, FOXA, C/EBPβ, ATF4, ZKSCAN3, E2F1, and others regulate the transcription of autophagy genes (reviewed in [13]). More recently, non-coding RNAs such as miRNA, lncRNA, circRNA have also been implicated in autophagy gene regulation [14]. These studies point to the essential role of autophagy gene transcription in modulating the pathway. However, the focus of these reports is limited to individual regulatory mechanisms in isolated cell types, and a detailed study identifying regulators of autophagy genes is lacking. This lacuna is starker in the case of embryonic stem cells (ESCs) which have the unique property of self-renewal and the ability to differentiate into three germ layers. Hence, these cells require an efficient homeostatic system, and precisely regulated cellular and metabolic turnover pathways such as autophagy [15]. While reports emphasize the importance of balanced autophagy levels in governing the ESC state [16–18], how these levels are maintained in the context of transcriptional regulation of autophagy genes in steady-state and cell state transition is less understood. This is especially important as autophagy has been implicated as an essential determinant for tissue differentiation as well as the self-renewal and differentiation potential of embryonic and adult stem cells [19, 20]. It also has roles in vasculogenesis, angiogenesis, skeletal muscle maintenance, regeneration, and immune system homeostasis [21]. Furthermore, embryonic and adult development and cell differentiation are associated with cell death, with significant cross-talk existing between autophagy and cell death as well as the genes involved in both pathways [22]. In fact, autophagic cell death is crucial for normal development in some organisms [23].

Determining whether changes in autophagy during stem cell differentiation correlate with changes in autophagy gene expression, and dissecting the mechanisms regulating these genes, would help open up potential avenues to identify druggable targets for autophagy modulation. This becomes pertinent given the background of mis-regulated autophagy genes being implicated in degenerative disorders, tumorigenesis and cancer progression, genome instability, impaired immune signalling and pathogen degradation among other disorders [24–26]. This study focuses on understanding the regulation of autophagy in mESCs in steady-state and during the early stages of differentiation. We observe an induction of autophagy as mESCs differentiate, which is accompanied by the upregulation of a subset of autophagy genes. Using a small molecule epigenetic inhibitor screen, we identify chromatin modifiers such as BRD4, G9A, EED and EZH2 as repressors of autophagy in mESCs. We further determine a novel role for EZH2 in autophagy maintenance during mESC differentiation, by regulating the expression of the autophagy gene, *Dram1*. Inhibition of EZH2 leads to the derepression of *Dram1*, which correlates with a shift from autophagy induction to caspase-independent cell death in differentiating mESCs. Thus, epigenetic regulation of *Dram1* by EZH2 helps to maintain the autophagy- cell death balance during mESC differentiation. The data shown in this study point to the crucial role of chromatin-mediated regulation of autophagy genes in governing this pathway as cells differentiate, and open new avenues for using epigenetic inhibitors, as druggable solutions for autophagy-mediated disorders.

## Materials and methods

### Cell culture and differentiation

V6.5 mESCs were cultured in the absence of feeders on 0.2% gelatin-coated plastic tissue culture dishes (Corning). Cells were grown in Knockout DMEM (Gibco, 10829-018) containing 15% FBS (Gibco, 10270-106), 2mM L glutamate 100X (Gibco, 25030-081), 1X penicillin/streptomycin (Gibco, 15140- 122), 1mM 100X MEM non-essential amino acids (Gibco, 11140-050), 2-mercaptoethanol 1000X (Gibco, 21985-023) and 1000U LIF (Leukaemia Inhibitory Factor) prepared in-house. For neural differentiation in monolayer culture, cells were grown in -LIF media containing 500nM retinoic acid. Media was changed every day, and cells were harvested at different time points. For autophagy induction, cells were treated with 2.5mM Rapamycin (ThermoFisher, PHZ1235) for 12 hours or 7.5μM Chloroquine (Sigma, C6628) for 8 hours. For autophagy inhibition, cells were treated with 4μM Wortmannin (Sigma, W1628) for 6 hours. For caspase inhibition, cells were treated with 20μM Z VAD-FMK (Sigma, 219007) for 3 hours.

### Establishment of mCherry-GFP-LC3 mESCs

The FUW mCherry-GFP-LC3 plasmid [27] was a gift from Anne Brunet (Addgene, 110060) and was co-transfected with psPAX2 (Addgene, 12260) and pMD2.G (Addgene, 12259) in 60% confluent HEK293T cells using FuGENE HD (Promega, E2311). 48 hours post transfection, viral supernatants were harvested and used for infection in the presence of 5μg/ml polybrene in V6.5 mESCs. The infected mESCs were subjected to FACS sorting and cells expressing high levels of GFP and mCherry were maintained as a polyclonal population and used in experiments. The cells were periodically checked for mycoplasma contamination.

### Epigenetic inhibitor treatment

Small molecule inhibitors for epigenetic pathways (Table S1) were a kind gift from the Structural Genomics Consortium. The drugs and their inactive analogs (used as negative controls) were dissolved in DMSO and working concentrations and treatment durations were optimized using standard protocols. The final treatment conditions are described in (Table S1).

### Autophagy detection

Cyto-ID (Enzo, ENZ-51031-K200) autophagy detection kit was used to assess autophagy levels in cells using published protocols [28].

### Confocal microscopy

The mCherry-GFP-LC3 mESCs were used to detect autophagy using confocal microscopy with a Nikon Eclipse Ti2 microscope using the 64X objective. The GFP and mCherry puncta were quantified using FIJI (ImageJ). After removal of background and setting appropriate intensity thresholds, the number of GFP and mCherry puncta were counted, and puncta per cell were calculated using DAPI as nuclear marker.

### Immunoblot

Total proteins were extracted from cells using RIPA buffer containing proteinase inhibitors on ice, followed by centrifugation at 12,000 rpm for 20 min at 4 °C. Protein concentration was measured using Bradford’s reagent. Total protein was subjected to SDS-PAGE under reducing conditions followed by transfer on to a PVDF membrane. The membrane was blocked using 5% BSA in Tris-buffered saline (TBS). Post blocking, the membrane was incubated at 4 °C overnight with the appropriate primary antibody. After 3 × 10 min wash in 1X TBS containing 0.1% Tween-20 (TBS-T), the membranes were incubated with an HRP-conjugated secondary antibody for 1 hour at room temperature. The membranes were developed using Femto substrate development kit (Invitrogen, 34094) and images were captured using a chemi-doc system (GE Healthcare, AI600). The following primary and secondary antibodies were used in the study: αTubulin (Sigma, T5168): 1:5000; LC3 (ProteinTech, 12135-1-AP): 1:1000; H3K27me3 (CST, 9733T): 1:1000; Caspase 3 (CST, 9662S): 1:1000; Anti rabbit HRP (Invitrogen, G-21234): 1:2000; Anti mouse HRP (Invitrogen, G21040): 1:2000.

### qRT PCR

Trizol (Life Technologies,15596018) was used to isolate total RNA from cells. After DNAse treatment, RNA was quantified using a NanoDrop spectrophotometer. Verso cDNA synthesis kit (Thermo Scientific, AB-1453) was used to generate cDNA. Diluted cDNA was used for qRT-PCR using gene specific primers (Table S2) and Power SYBR Green PCR Master Mix (Applied Biosystems, A25742). *Gapdh* was used as an endogenous control.

### *Dram1* siRNA generation

Template for esiRNA [29, 30] production against specific genes was prepared by PCR amplification from mESC or MEF cDNA. *In vitro* transcription was performed using T7 RNA polymerase, followed by digestion of the double-stranded RNA using RNase III. RNA was then transfected into mESCs using the DharmaFECT 1 transfection reagent (Dharmacon, T-2001-02). After 72 hours, cells were lysed in TRIzol for RNA, or in RIPA lysis buffer to prepare whole-cell protein extract. Non-targeting esiRNA was prepared using GFP as a template in all experiments.

### Chromatin immunoprecipitation (ChIP)

mESCs were grown in 10 cm plates at 70% confluency. After trypsinization, cells were fixed for 10 min at 37 °C using a final concentration of 1% formaldehyde. Glycine was added to a final concentration of 125mM to quench the crosslinking reaction. Cells were washed twice with ice cold 1x PBS. 1 ml SDS lysis buffer was added to each sample and incubated at 4 degrees for 30 minutes. This was followed by sonication using a BioruptorTM (UCD200) at high setting for 30 seconds ON, 60 seconds OFF for 25 cycles. Subsequent steps were performed as described in [31]. The purified DNA was analysed by qPCR using specific primers as described in Table S2. The antibodies used for ChIP are as follows: H3K27me3 (Invitrogen, MA5-11198): 4μg. H3K4me3 (Diagenode, C15410003-50): 2μg.

## Results

### Autophagy is induced during RA-mediated mESC differentiation

To determine autophagy status in mESCs, we established a stable mESC line expressing the autophagosome membrane protein LC3, tagged with GFP and mCherry (henceforth called LC3-tagged- mESCs) [27]. Autophagosomes were identified by the presence of GFP as well as mCherry positive puncta, while autophagic flux was detected by the loss of GFP positive puncta as autophagosomes fuse with the lysosomes. Untreated mESCs showed fewer and diffused GFP and mCherry positive puncta (Figure S1a), while induction of autophagy by the known autophagy activator rapamycin, led to an increase in both GFP and mCherry positive puncta indicating an increase in both autophagy initiation and flux. Treatment by the autophagosome-lysosome fusion inhibitor chloroquine however, led to an increase in GFP positive puncta and a reduction of mCherry positive puncta, consistent with the accumulation of autophagosomes and block in autophagic flux (Figure S1a and S1b). These cells therefore, were reliable indicators of autophagy levels in mESCs at steady state as well as in response to autophagy modulators. We further treated these cells with retinoic acid (RA) to determine the status of autophagy during mESC differentiation along the neural lineage. The cells expressed neural markers such as *Nestin* and *βIII tubulin* and lost pluripotency markers such as *Oct4* and *Nanog* after 3 days of RA treatment (Figure S1c). Undifferentiated mESCs exhibited largely diffused GFP and mCherry, with few GFP-positive puncta and a higher number of mCherry puncta, consistent with basal levels of autophagy in these cells. RA treatment however, resulted in the increased formation of distinct GFP and mCherry positive puncta indicating an increase in autophagy initiation and autophagic flux (Figure 1a and 1b). We verified this increase in autophagy at the protein level by immuno-blotting with an antibody against LC3. We observed an increase in LC3 II levels and LC3 II/LC3 I ratio upon RA treatment of mESCs indicating an induction of autophagy (Figure 1c and 1d).

**Figure 1:**
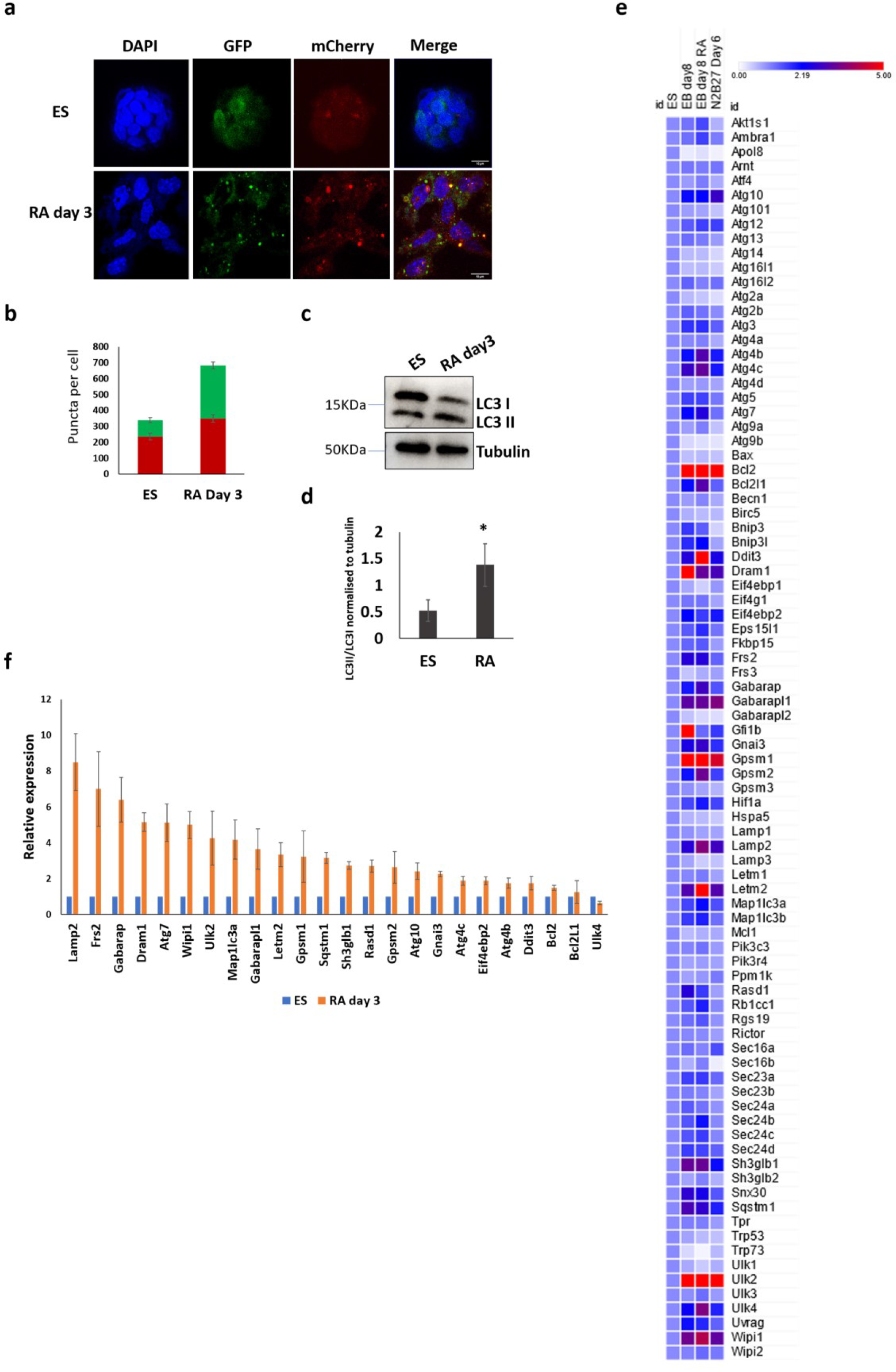
Autophagy is induced during RA-mediated differentiation of mESCs. 1a: GFP and mCherry positive puncta in LC3-tagged-ES cells (Top panel), and RA-treated cells (Bottom panel). Nuclei are labelled with DAPI. Scale bar =10μm. 1b: Quantitation of GFP and mCherry positive puncta in mESCs and RA-treated cells. Y axis represents average puncta per cell. N=150 cells. Error bar indicates standard error of the mean (SEM). 1c: Immunoblot depicting LC3I and LC3II in mESCs and RA-treated cells. α tubulin is used as loading control. 1d: Quantitation of immunoblot in mESCs and RA- treated cells. After quantification of band intensity, LC3II/LC3I levels were normalised to α tubulin levels. N=3 biological replicates. Asterisks indicate statistically significant differences compared to ES (*: p< 0.05, paired t-test). 1e: Gene expression analysis of autophagy genes in mESCs, EB day 8 (ES medium-LIF), EB day 8 (RA) and RA treated cells day 6 (N2B27 medium, monolayer). Scale bar represents fold change. Data is presented as a heatmap depicting average fold change normalised to mESCs. The heatmap was plotted using Morpheus, https://software.broadinstitute.org/morpheus 1f: RT-qPCR analysis of autophagy genes in mESCs and RA-treated cells. Y axis represents fold change wrt *Gapdh* normalised to mESCs. N=3 biological replicates. Error bar indicates standard error of the mean (SEM).

To determine whether the induction of autophagy during mESC differentiation was accompanied by an increase in the expression of autophagy genes, we analysed published RNA-Seq data (GSE84401 [32]) to determine the expression levels of autophagy genes in mESCs, embryoid bodies (EBs), RA treated EBs and RA treated mESCs grown in N2B27 media. We found that out of 86 genes, 24 genes were significantly upregulated (>2 fold) upon mESC differentiation (Figure 1e and S1d). Interestingly, the most highly upregulated genes were regulators of autophagy rather than genes coding for constituents of the pathway itself. These include autophagy activators such as *Dram1, Ulk2, Gfi1b and Wipi1*. Genes coding for antiapoptotic proteins such as BCL2 and GPSM1 were also upregulated in differentiated cells. We validated the upregulation of select genes in mESCs treated with RA by RT-qPCR. We saw a significant upregulation in the expression of genes such as *Lamp2, Gabarap, Ulk2, Dram1, Wipi1* among others (Figure 1f). Among the Atg genes, *Atg4b*, *Atg4c*, *Atg7*, and *Atg10* were upregulated in RA treated mESCs. Our studies indicate that autophagy is induced in RA-mediated differentiation of mESCs and this is associated with an increased expression of a subset of autophagy genes.

### Autophagy genes are regulated by epigenetic pathways

As mESC differentiation led to an upregulation of autophagy genes, we sought to identify transcriptional regulators of these genes. Epigenetic regulatory mechanisms play a crucial role in determining transcriptional outcomes in response to stimuli. To identify the epigenetic regulators of autophagy, we used small molecule inhibitors (Table S1) targeting specific epigenetic pathways. We used a fluorescence-based autophagy detection kit (Cyto-ID) to detect autophagy levels. We observed an increase in autophagy in mESCs treated with inhibitors such as GSK343 (EZH2), UNC1999 (EZH2), A-395 (EED), UNC0642(G9a/GLP), A366 (G9a/GLP), and JQ1 (BET domain proteins) (Figure 2a); while treatment with other inhibitors did not show a significant change in autophagy (Figure S2a). Treatment with chemically inactive analogues such as UNC2400 (EZH2-ve), A395N (EED -ve), -JQ1 (BET domain - ve) did not show significant changes in autophagy (Figure 2a), indicating specific effects of certain epigenetic inhibitors on autophagy. We validated the FACS data by treating LC3-tagged-ES cells with GSK343, A366, and JQ1; and observed an induction of autophagy as seen by an increase in GFP and mCherry positive puncta (Figure 2b and 2c). Our screen identified the Polycomb group proteins EED and EZH2 as regulators of autophagy in mESCs. Inhibition of EZH2 using GSK343 as well as UNC1999 led to a significant induction of autophagy (Figure 2a, 2b and 2c). FACS analysis in LC3-tagged-ES cells revealed an in both GFP and mCherry positive puncta upon treatment with GSK343. However, the increase of mCherry puncta was higher than that of the GFP puncta indicating an increase in autophagic flux upon EZH2 inhibition (Figure 2c). RT-qPCR analysis showed the mis-regulation of several autophagy genes upon treatment with small molecule epigenetic inhibitors (Figure 2d). Genes such as *Becn1, Lc3a and Ulk2* were upregulated upon treatment with all candidate inhibitors, while other genes responded in an inhibitor-dependent manner, indicating that different epigenetic molecules regulate the expression of distinct autophagy genes. Interestingly, genes such as *Lamp2, Dram1, and Ulk2* which were upregulated upon RA treatment (Figure 1e and 1f) were also upregulated upon inhibition of EZH2 as well as EED in mESCs (Figure 2d), indicating a putative role for the Polycomb group proteins in the regulation of these genes during mESC differentiation. Taken together, these results indicate that in mESCs, autophagy is repressed by epigenetic modulators such as G9a/GLP, EZH2, EED and BET proteins and inhibition of these pathways results in a concomitant upregulation of autophagy genes.

**Figure 2:**
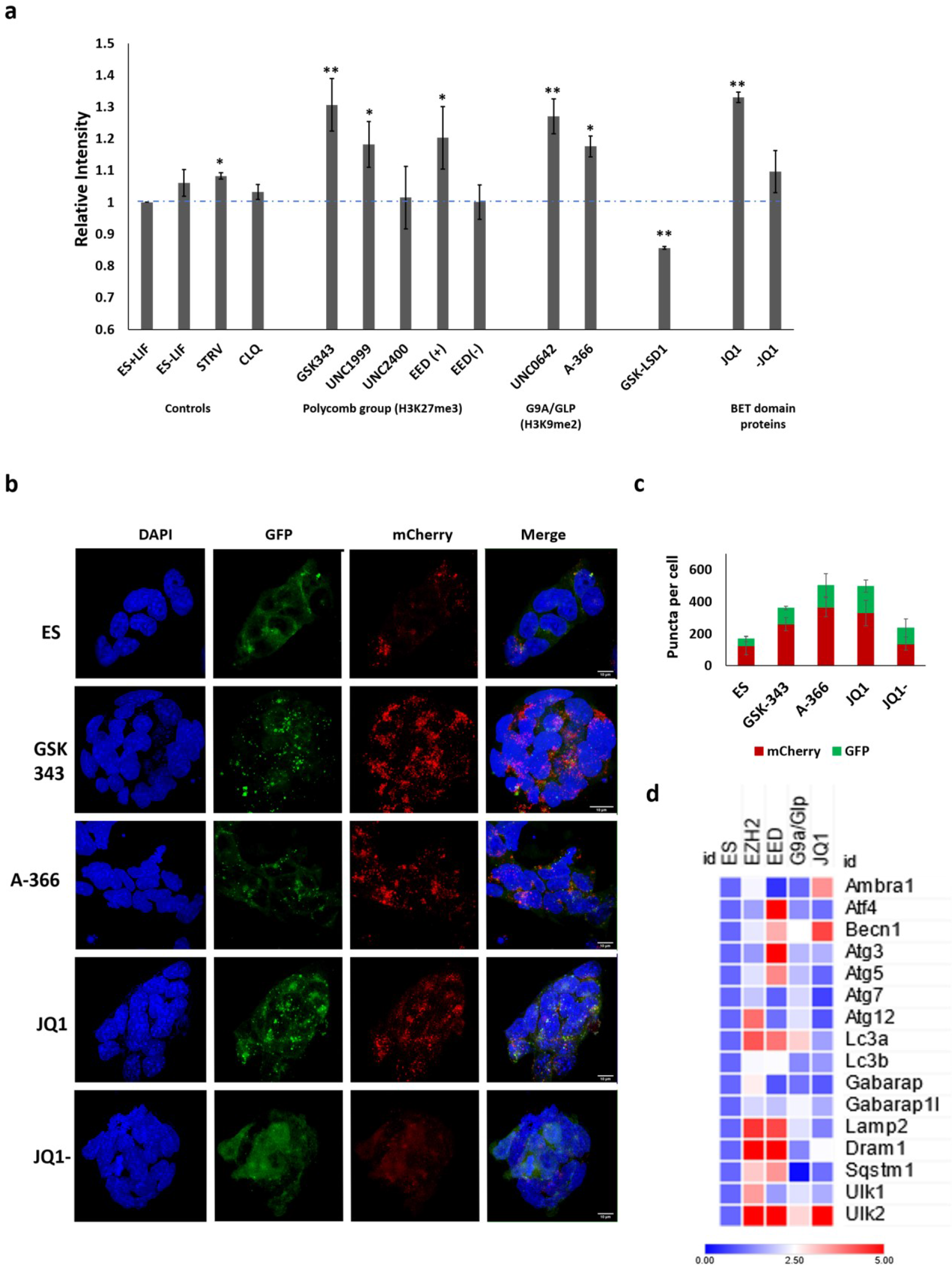
Autophagy is regulated by epigenetic pathways in mESCs. 2a: GFP intensity depicting autophagy levels in mESCs and cells treated with epigenetic inhibitors. Y axis depicts relative fluorescence intensity normalised to mESCs. N=3.biologocal replicates. Error bar indicates standard error of the mean (SEM). Asterisks indicate statistically significant differences compared to ES+LIF (*: p< 0.05, **: p<0.005; paired t-test). 2b: GFP and mCherry positive puncta in ES cells (Top panel) and cells treated with epigenetic inhibitors (Bottom panels). Nuclei are labelled with DAPI. Scale bar =10μm. 2c: Quantitation of GFP and mCherry puncta in mESCs and cells treated with epigenetic inhibitors. Y axis represents average puncta per cell. N=150 cells. 2d: Expression of autophagy genes in mESCs and cells treated with epigenetic inhibitors. Data is presented as a heatmap depicting average fold change normalised to mESCs. N=3. The heatmap was plotted using Morpheus, https://software.broadinstitute.org/morpheus.

### EZH2 inhibition leads to caspase-independent, autophagy-associated cell death in differentiating mESCs

Polycomb group proteins regulate numerous cellular pathways that play crucial roles in cellular homeostasis, development and differentiation, and the enzyme EZH2 is the primary determinant for the establishment of the repressive H3K27me3 modification of target sites. Given the specific roles of Polycomb proteins in stem cells and during neuronal differentiation [33–36], we focussed our further experiments on dissecting the role of the Polycomb enzyme EZH2 in autophagy regulation in mESCs and differentiation. Observation of cells under brightfield showed that while mESCs treated with GSK343 did not exhibit significant changes in morphology, RA-treated cells exposed to GSK343, showed extensive cell death (Figure S2b). Cell counting assay showed that compared to untreated mESCs, 49% of cells remained on day 3 of RA treatment and this number dropped to 8% when cells were treated with RA_GSK343 (Figure S2c). Treatment with GSK343 alone however, did not significantly affect the doubling of ES cells. While programmed cell death is a hallmark of neural differentiation in normal development, RA-mediated differentiation, as well as in neuroblastomas [37–40]; treatment with RA_GSK343 exacerbated this phenomenon and we found no viable cells beyond day 4 of treatment. We limited our further treatment duration to 48 hours as extensive cell death was seen beyond this point. To determine whether the cell death phenomenon was apoptosis-dependent, we performed Annexin-PI staining followed by FACS analysis. We observed that, consistent with previous reports [37], there was an increase in the percentage of Annexin-positive (Early apoptotic) and Annexin+PI-positive (Late apoptotic) cells upon RA treatment of mESCs. In RA_GSK343 treated cells however, there was a significant increase in PI-positive/Annexin-negative as well as Annexin+PI-positive cells, while the percentage of only Annexin-positive cells remained largely unchanged (Figure 3a and S2d). The increase in PI-positive/Annexin-negative cells in RA_GSK343 treated cells indicates an induction of non-apoptotic cell death. To determine whether the cell death was Caspase-dependent, we performed immunoblotting with a Caspase-3 antibody and saw no change in cleaved Caspase-3 levels (Figure 3b) in RA_GSK343 treated cells. We validated this by testing the effect of the pan-Caspase inhibitor, Z VAD-FMK, on RA_GSK343 treated cells, with etoposide treated ES cells as a positive control for cell death. While the percentage of Annexin-positive and Annexin+PI-positive cells were reduced in Etoposide+Caspase inhibitor treated cells, we observed no significant changes in the presence of RA_GSK343+Caspase inhibitor (Figure S2e), further confirming that the cell death phenotype is Caspase independent. RT-qPCR using primers for pro-apoptotic genes also revealed no significant changes at the transcript level (Figure S2f).

**Figure 3:**
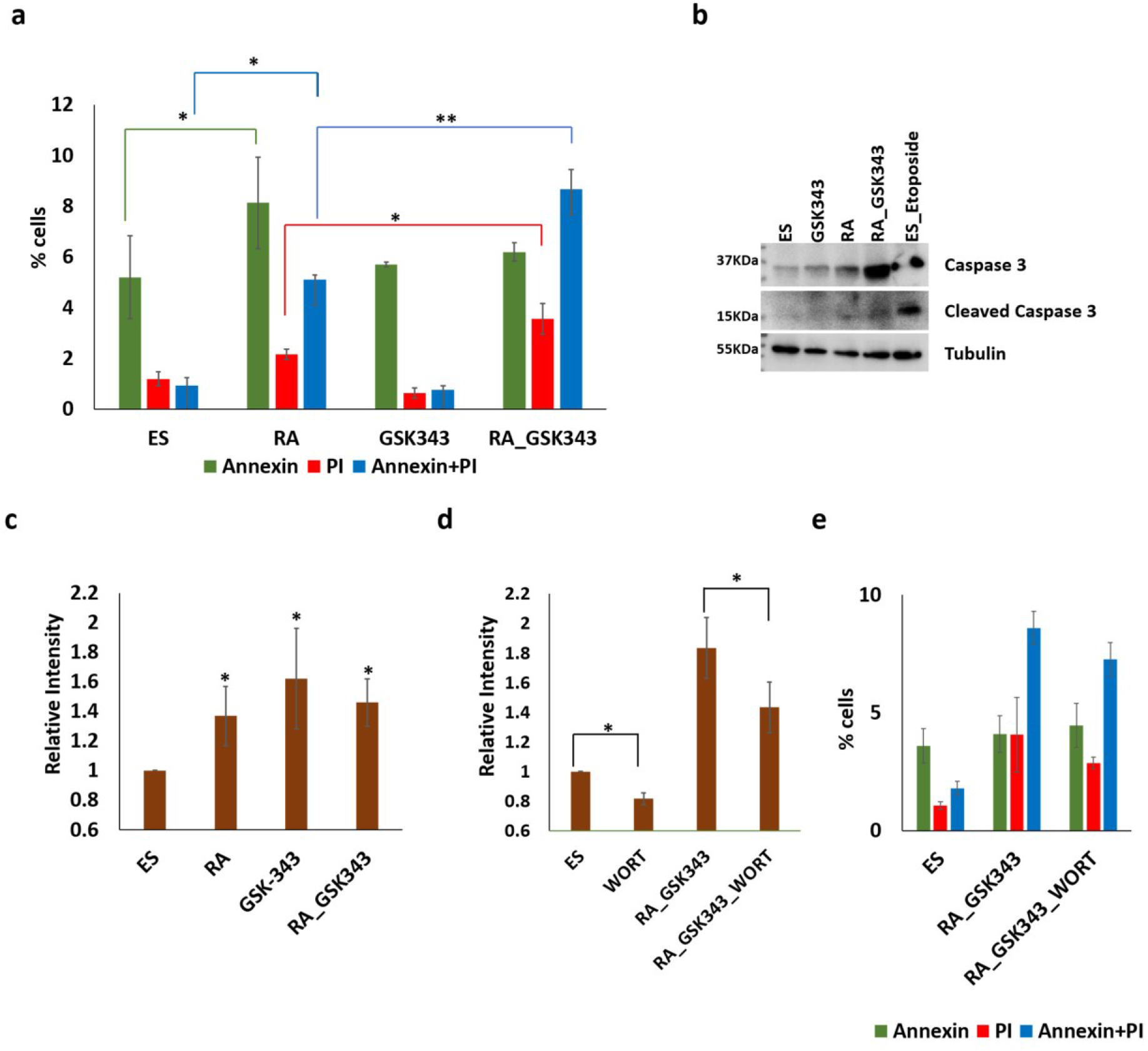
GSK343 treatment leads to autophagy-associated, caspase-independent cell death in differentiating mESCs. 3a: Annexin-PI assay in mESCs; RA, GSK343 and RA_343 treated cells. Y axis represents percentage of Annexin positive, PI-positive and Annexin+PI positive cells. N= 3 biological replicates. Error bar indicates standard error of the mean (SEM). Asterisks indicate statistically significant differences (*: p< 0.05, **: p<0.005; paired t-test). 3b: Immunoblot depicting intact caspase 3 and cleaved caspase 3 in mESCs; RA, GSK343 and RA_343 treated cells. mESCs treated with etoposide were used as positive control for caspase-dependent apoptosis. α-tubulin is used as loading control. N= 3 biological replicates. 3c: GFP intensity depicting autophagy levels in mESCs; RA, GSK343 and RA_GSK343 treated cells. Y axis depicts relative fluorescence intensity normalised to mESCs. Error bar indicates standard error of the mean (SEM). Asterisks indicate statistically significant differences compared to ES (*: p< 0.05, **: p<0.005; paired t-test). N= 3 biological replicates. 3d: GFP intensity depicting autophagy levels in mESCs, Wortmannin treated mESCs, RA_GSK343 treated cells and RA_GSK343+ Wortmannin treated cells. Y axis depicts relative fluorescence intensity normalised to mESCs. N= 3 biological replicates. Error bar indicates standard error of the mean (SEM). Asterisks indicate statistically significant differences (*: p< 0.05; paired t-test). 3e: Annexin-PI assay in mESCs, RA_GSK343 and RA_GSK343+Wortmannin treated cells. Y axis represents percentage of Annexin-positive, PI-positive and Annexin+PI-positive cells. N= 3 biological replicates.

Autophagic cell death (Type II cell death) refers to non-apoptotic cell death that is accompanied by increase in autophagy [41]. Because the causative relationship between autophagy and cell death is difficult to assess [42], the guidelines for the nomenclature of autophagic processes suggest that type II cell death can be classified as autophagy-mediated or autophagy-associated depending on whether the observed cell death is accompanied by an increase in autophagy and whether chemical or genetic inhibition of autophagy is able to rescue cell death [43, 44]. To determine whether EZH2 inhibition in differentiating mESCs caused autophagic cell death, we assessed the levels of autophagy in RA_GSK343 treated cells. We observed that treatment with RA, GSK343 and RA_GSK343 led to a robust induction of autophagy, however, RA_GSK343 treatment led to similar levels of induction as individual treatments (Figure 3c and S3a). This may indicate that the processes by which GSK343 and RA activate autophagy, may be overlapping. The overlap between autophagy genes upregulated in RA as well as GSK343 treated cells (Figures 1e,1f and 2d) also points to a potential common mechanism of autophagy activation by the two treatment conditions. To determine whether the cell death phenotype was autophagy-dependent, we treated the cells with autophagy inhibitor Wortmannin. Wortmannin treatment led to a reduction in autophagy in RA_GSK343 treated cells (Figure 3d). However, Annexin-PI staining showed a modest, statistically insignificant reduction in cell death upon wortmannin treatment (Figure 3e). This indicates that while the cell death seen in RA_GSK343 treated cells is accompanied by an increase in autophagy, it may not be completely autophagy-dependent.

### EZH2 targets *Dram1* to regulate autophagy and cell death

As treatment with RA and GSK343 led to an increase in autophagy and RA_GSK343 treatment led to an increase in autophagy along with cell death, we sought to dissect the possible mechanism that regulated this phenotype. GSK343 is a selective inhibitor of the Polycomb group enzyme EZH2, that establishes the H3K27me3 mark to repress target genes [45, 46]. To determine whether the induction of autophagy and cell death in RA_GSK343 treated cells was associated with changes in autophagy gene expression, we compared the genes upregulated in RA and GSK343 treatment (Figures 1e,1f and 2d). We identified genes such as *Lamp2, Gabarap and Dram1*, upregulated in RA as well as GSK343 treated cells. The gene coding for the DNA damage-regulated autophagy modulator 1 (DRAM1) was of special interest, as this protein is shown to activate autophagy as well as cell death [47–49] and could potentially serve as a regulator of both phenotypes observed in our experiments. *Dram1* was upregulated in mESCs treated with RA as well as GSK343; however, RA_GSK343 treatment led to a further compounded upregulation of *Dram1*, many times higher than individual treatments (Figure 4a). We used an siRNA-mediated *Dram1* knockdown approach to determine the role of *Dram1* in the induction of autophagy and cell death. We saw a significant decrease in *Dram1* expression in siRNA treated cells (*Dram1* KD) (Figure S3b). We then characterised the effect of *Dram1* knockdown on autophagy and cell death levels in mESCs as well as differentiating cells. CytoID assay revealed that autophagy levels remained unchanged in control, RA and RA_343 treated *Dram1* KD cells, however, significant reduction in autophagy was seen in GSK343 treated mESCs (Figure 4b). This indicates that the autophagy induction seen in RA treated cells may not depend on *Dram1*. This data also suggests that while autophagy induction is seen in RA_GSK343 treated cells, knockdown of *Dram1* is insufficient to reverse this induction and may point to other regulatory modules. Other upstream autophagy regulators such as *Gabarap* and *Lamp2* which were also upregulated in RA treated cells may contribute to the induction of autophagy.

**Figure 4:**
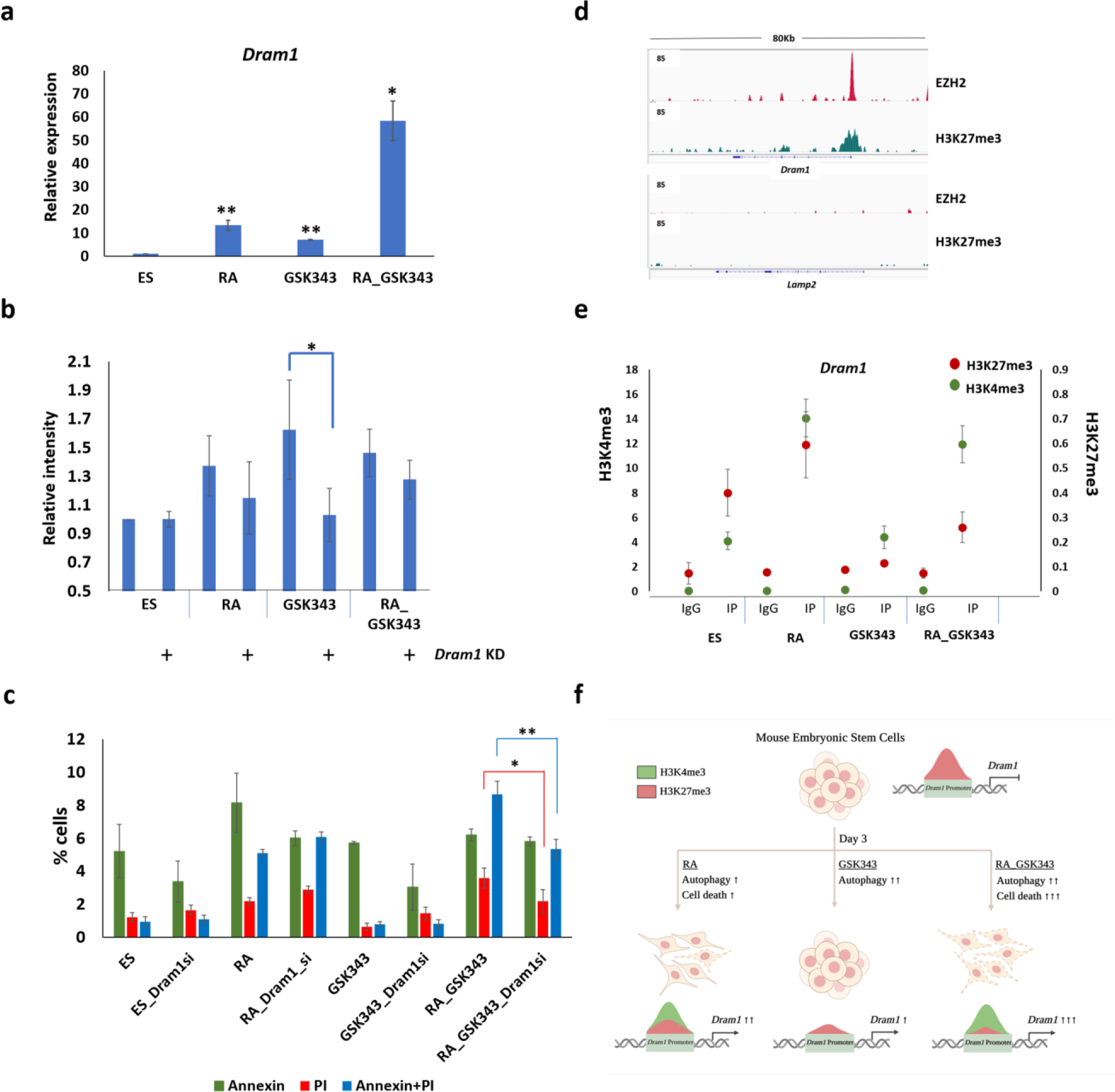
Expression of the autophagy gene *Dram1* is regulated by EZH2 in mESCs and differentiating cells. 4a: RT-qPCR analysis of *Dram1* expression in mESCs; RA, GSK343 and RA_GSK343 treated cells. Y axis represents fold change wrt *Gapdh* normalised to mESCs. N=3 biological replicates. Error bar indicates standard error of the mean (SEM). Asterisks indicate statistically significant differences compared to mESCs (*: p< 0.05, **: p<0.005; paired t-test). 4b: GFP intensity depicting autophagy levels in mESCs; RA, GSK343 and RA_GSK343 treated cells without and with *Dram1* siRNA. Y axis depicts relative fluorescence intensity normalised to ES cells. N= 3 biological replicates. Error bar indicates standard error of the mean (SEM). Asterisks indicate statistically significant differences. (*: p< 0.05, paired t-test). 4c: Annexin-PI assay in mESCs, RA, GSK343 and RA_GSK343 treated cells without and with *Dram1* siRNA. Y axis represents percentage of Annexin-positive, PI-positive and Annexin+PI positive cells. N= 3 biological replicates. Error bar indicates standard error of the mean (SEM). Asterisks indicate statistically significant differences (*: p< 0.05, **: p<0.005; paired t-test). 4d: EZH2 and H3K27me3 enrichment over *Dram1* (Top panel) and *Lamp2* (Bottom panel). IGV Snapshot of ChIP-Seq enrichment is depicted. 4e: ChIP-qPCR analysis depicting the enrichment of H3K4me and H3K27me3 at the *Dram1* promoter. The Y axes represent percentage of input. Error bar indicates standard error of the mean (SEM). 4f: Ezh2 regulates *Dram1* expression to maintain autophagy-apoptosis balance in differentiating ES cells. In mESCs, *Dram1* is repressed by the establishment of H3K7me3 by the Polycomb enzyme EZH2. During RA mediated differentiation, *Dram1* is upregulated and the activating H3K4me3 mark is established on the promoter. However, the H3K27me3 mark is retained on the promoter, leading to a bivalent chromatin domain. This is accompanied by an increase in autophagy and cell death, with cell death levels restricted to 30-50%. In the presence of the inhibitor for Polycomb protein EZH2, H3K27me3 levels over the *Dram1* promoter are drastically reduced. This, coupled with the activating H3K4me3 mark, leads to “hyper-expression” of *Dram1*. This results in a shift between the autophagy and cell death balance and results in a drastic increase in caspase-independent cell death. Thus, a Polycomb mediated regulation of Dram1 levels during ES cell differentiation maintains the balance between autophagy and cell death.

To determine the effect of Dram1 depletion on cell death, we performed Annexin-PI staining. Interestingly, we observed that *Dram1* KD was able to partially rescue cell death only in RA_GSK343 treated cells, where we detected significant reduction of Annexin-negative/PI-positive as well as Annexin+PI-positive cells (Figure 4c). This effect was not seen in other treatment conditions. Our results point to distinct effects of *Dram1* KD on autophagy and cell death in mESCs and differentiating cells which may indicate that *Dram1* plays a cell state specific role in regulating autophagy vs. cell death. This regulation may be mediated by the Polycomb complex, as *Dram1* upregulation by the EZH2 inhibitor primarily affected autophagy in mESCs, and cell death in differentiating cells.

*Dram1* has been identified as a p53 dependent regulator of autophagy and apoptosis [48]. To determine whether the effect of GSK343 on *Dram1* is mediated by p53, we determined the levels of p53 transcript in our cells. We observed no significant changes in p53 in any of our treatments (Figure S3c). This indicates that *Dram1* transcription may directly be regulated by EZH2. To determine whether EZH2 is enriched at the *Dram1* promoter, we analysed published ChIP-Seq data (GSE89929) [50], and observed an enrichment of EZH2 as well as the H3K27me3 mark at the *Dram1* promoter in mESCs. In contrast, *Lamp2*, which was also upregulated in RA treated cells remained devoid of both EZH2 and H3K27me3 enrichment (Figure 4d). This indicates that the *Dram1* promoter is selectively targeted by EZH2. The upregulation of *Lamp2* upon GSK343 treatment may be attributed to an indirect mechanism involving upstream regulators of the gene. As our studies indicated that *Dram1* upregulation leads to distinct effects on autophagy and cell death in mESCs vs differentiating cells, we analysed the enrichment of activating and repressive chromatin modifications on the *Dram1* promoter in these cells. ChIP-Seq data (GSE135318) [51], revealed that the *Dram1* promoter is enriched with primarily the H3K27me3 mark in mESCs. In neural progenitor cells (NPCs) however, we observed the enrichment of H3K27me3 as well as H3K4me3 at the *Dram1* promoter (Figure S3d). The increase in the active mark H3K4me3 in NPCs is consistent with the increased expression of *Dram1*. This pattern of chromatin modifications, called the bivalent domain is commonly present on promoters of genes that are repressed but poised for activation in stem cells [52]. We validated this in mESCs and RA treated cells by ChIP-qPCR and observed that while the *Dram1* promoter was marked with H3K27me3 in mESCs, a bivalent domain (H3K27me3 and H3K4me3) is seen in RA treated cells. Upon GSK343 treatment, H3K27me3 is reduced, leaving high enrichment of H3K4me3 in RA_GSK343 cells (Figure 4e and S3e), which correlates with the compounded increase in *Dram1* expression. Taken together, our results indicate that in mESCs, the *Dram1* gene is repressed by the action of Polycomb group proteins and the deposition of H3K27me3 on the promoter. As cells differentiate, activating marks such as H3K4me3 are deposited, which contribute to the upregulation of *Dram1* and induction of autophagy. However, the H3K27me3 mark is retained on the promoter resulting in the establishment of a bivalent domain, which possibly limits the expression of *Dram1*. Treatment of differentiating cells with the EZH2 inhibitor GSK343 abolishes the H3K27me3 mark, and this coupled with the activating H3K4me3 modification, results in a “hyper-expression” of Dram1 and extensive cell death (Figure 4f). Thus, Polycomb mediated regulation of the *Dram1* gene helps to regulate autophagy in mESCs and cell death during early RA-mediated differentiation.

## Discussion

This study presents a systematic analysis of autophagy regulation in mESCs and early neural differentiation. The increase in autophagy seen during RA-mediated differentiation of mESCs is consistent with previous studies that show autophagy induction during differentiation in the mouse olfactory bulb and also in RA treated mouse neuroblastoma cells [53, 54]. Several autophagy genes were upregulated in differentiating cells. Interestingly, the gene coding for the lysosomal protein LAMP2 was the one of most upregulated autophagy gene upon mESC differentiation, which is consistent with previous reports [18]. GABARAP has been shown to induce autophagy via ULK1 activation [55]. DRAM1 regulates autophagic flux by enhancing lysosomal acidification in an mTOR-dependent pathway [47, 56]. Our results are in line with previous reports which show that autophagy is induced during the differentiation of human ESCs, neural and cardiac stem cells [53,57,58]. However, autophagy is reported to be reduced during the differentiation of haematopoietic, dermal and epidermal stem cells [59]. The difference in trends between different stem cell types point to a context-specific role of autophagy in self-renewal and differentiation [59, 60]. It would be interesting to determine autophagy levels in mESCs subject to directed differentiation along different lineages. Targeted knock-out and overexpression experiments using the autophagy genes up-regulated in RA treated cells would help elucidate the causative effect of autophagy induction in mESC differentiation.

The small molecule epigenetic inhibitor screen identified G9A/GLP, EZH2 and BET domain proteins as autophagy repressors in mESCs. G9A/GLP has been reported as a repressor of autophagy in HeLa cells, MEFs, and various cancer cell lines [10,61,62,62]. In contrast, *Drosophila* G9A activates starvation induced autophagy by activating the autophagy gene *Atg8a* [63], indicating a context specific role of the enzyme. It would be interesting to compare targets of G9A/GLP in different cell/model systems to determine the downstream effectors of autophagy regulation. The role of JQ1 as an activator of autophagy has been reported in studies on cancer cell lines as well as in viral infections and recovery after spinal cord injuries [64–67]. Our data is consistent with the role of JQ1 as an activator of autophagy in mESCs. Our study presents the Polycomb group proteins EZH2 and EED as novel regulators of autophagy in mESCs and indicates that EZH2 targets *Dram1* to regulate its expression. Our data also suggest that RA mediated differentiation of ES cells leads to an induction of autophagy as well as apoptosis. However, treatment with EZH2 inhibitor in differentiating cells leads to severe cell death which is independent of the caspase pathway. This function, while being associated with autophagy, may not be dependent on autophagy induction, as inhibition of autophagy was unable to rescue the severe cell death phenotype. It is tempting to hypothesize that the balance between autophagy and cell death is maintained in stem cells and early differentiation by regulating *Dram1* expression levels which in turn are governed by epigenetic pathways. The mechanism causing the shift between autophagy and cell death remains to be understood. DRAM1 is a lysosomal protein that regulates cell death by increasing the levels and lysosomal localisation of the pro-apoptotic protein BAX in a transcription independent manner [49]. While our results did not show a change in the transcript levels of pro-apoptotic genes in RA_GSK343 treated cells (Figure S2e), *Dram1* upregulation may lead to increase in BAX localization to lysosome which inhibits its degradation and enhances cell death. Additionally, while the focus of our study has been on the regulation of *Dram1* by EZH2, one cannot discount the possibility that GSK343 treatment would cause global transcriptional changes, indirectly affecting our phenotype. Polycomb group proteins perform regulatory functions in a myriad of cellular pathways [46] and the cell death phenotype may be a result of alternative regulatory pathways. ChIP-Seq and RNA-Seq experiments using RA_GSK343 treated cells would shed light on the genome-wide effects of the inhibitor treatment on EZH2 occupancy and gene expression; and enable an accurate dissection of the balance between autophagy and cell death.

Our study identifies small molecule epigenetic drugs that activate autophagy in mESCs. Epigenetic inhibitors have emerged as promising therapeutic alternatives to cytotoxic drug regimens [68]. Thus the identification of novel regulatory mechanisms for cytoprotective pathways such as autophagy not only provides an enhanced understanding of the role of autophagy in stemness and differentiation, but also facilitates new avenues into developing druggable treatment paradigms to combat autophagy-mediated disorders.

## Author contributions

D.P conceptualized, designed and performed the experiments of the project. A.K contributed to the experiments depicted in figures 1c,3b and 4a. G.S.B contributed to the experiments depicted in figures 1f,2d and 4d. D.P wrote the original draft. The work was carried out in D.S lab using grants awarded to D.P and D.S

## Supplemental Figures

**Supplemental Figure 1:**
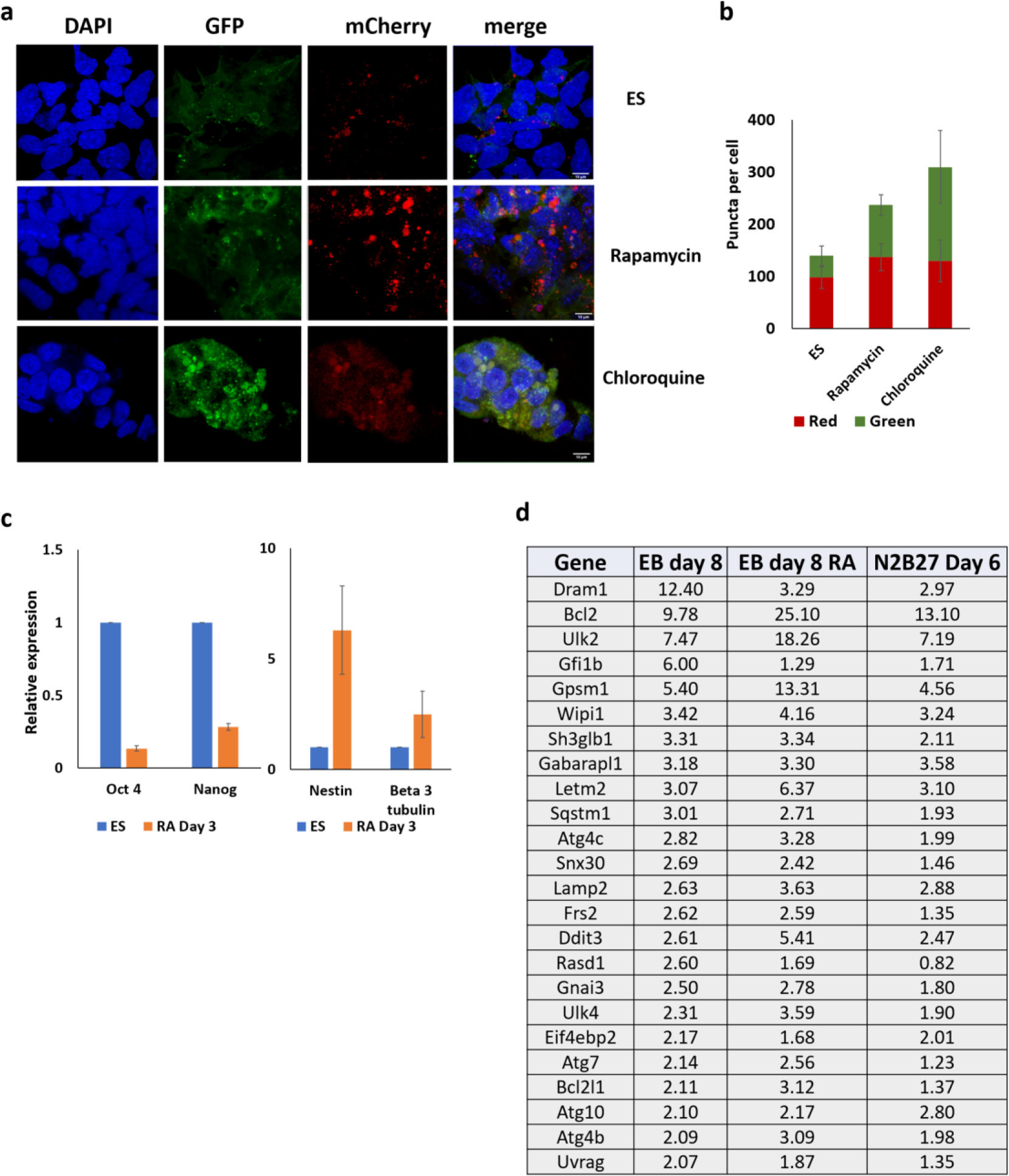
S1a: GFP and mCherry positive puncta in LC3-tagged mESCs and mESCs treated with Rapamycin and chloroquine. Nuclei are labelled with DAPI. Scale bar =10μm. S1b: Quantitation of GFP and mCherry positive puncta in LC3-tagged mESCs and mESCs treated with Rapamycin and chloroquine. Y axis represents average puncta per cell. N=150 cells. S1c: RT-qPCR analysis of pluripotency and neural gene expression in mESCs and RA-treated cells. Y axis represents fold change wrt *Gapdh* normalised to mESCs. N=3 biological replicates. S1d: List of autophagy genes upregulated upon neural differentiation of mESCs. Fold change normalised to mESCs is depicted.

**Supplemental figure S2:**
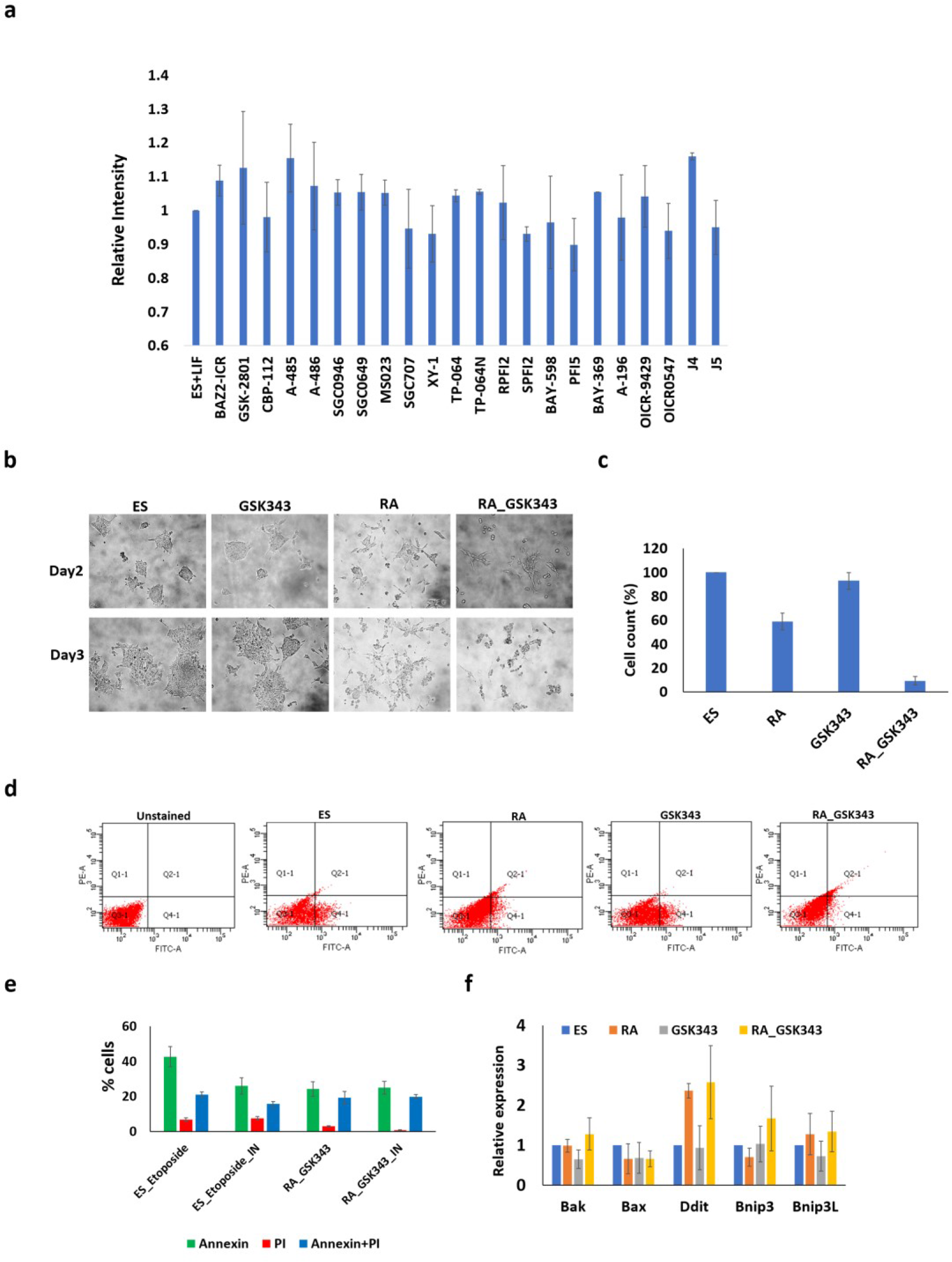
S2a: GFP intensity depicting autophagy levels in mESCs and cells treated with epigenetic inhibitors. Y axis depicts relative fluorescence intensity normalised to mESCs. S2b: Brightfield images depicting the morphology of mESCs, RA, GSK343 and RA_GSK343 treated cells. Images for day 2 and day 3 of treatment are shown. S2c: Cell counting assay depicting cell numbers after 3 days of plating equal numbers of mESCs, RA, GSK343 and RA_GSK343 treated cells. The Y axis represents % of cells normalised to mESCs. S2d: Scatter plot depicting FACS analysis of Annexin V-FITC and PI-PE positive cells in mESCs, RA, GSK343 and RA_GSK343 treated cells. Unstained cells were used as control. S2e: Annexin-PI assay in Etoposide treated ES cells, and RA_343 treated cells in the absence and presence of Caspase inhibitor. Y axis represents percentage of Annexin positive, PI-positive and Annexin+PI positive cells. N= 3 biological replicates. S2f: RT-qPCR analysis of pro-apoptotic gene expression in mESCs, RA, GSK343 and RA_GSK343 treated cells. Y axis represents fold change wrt *Gapdh* normalised to mESCs. N=3.

**Supplemental figure S3:**
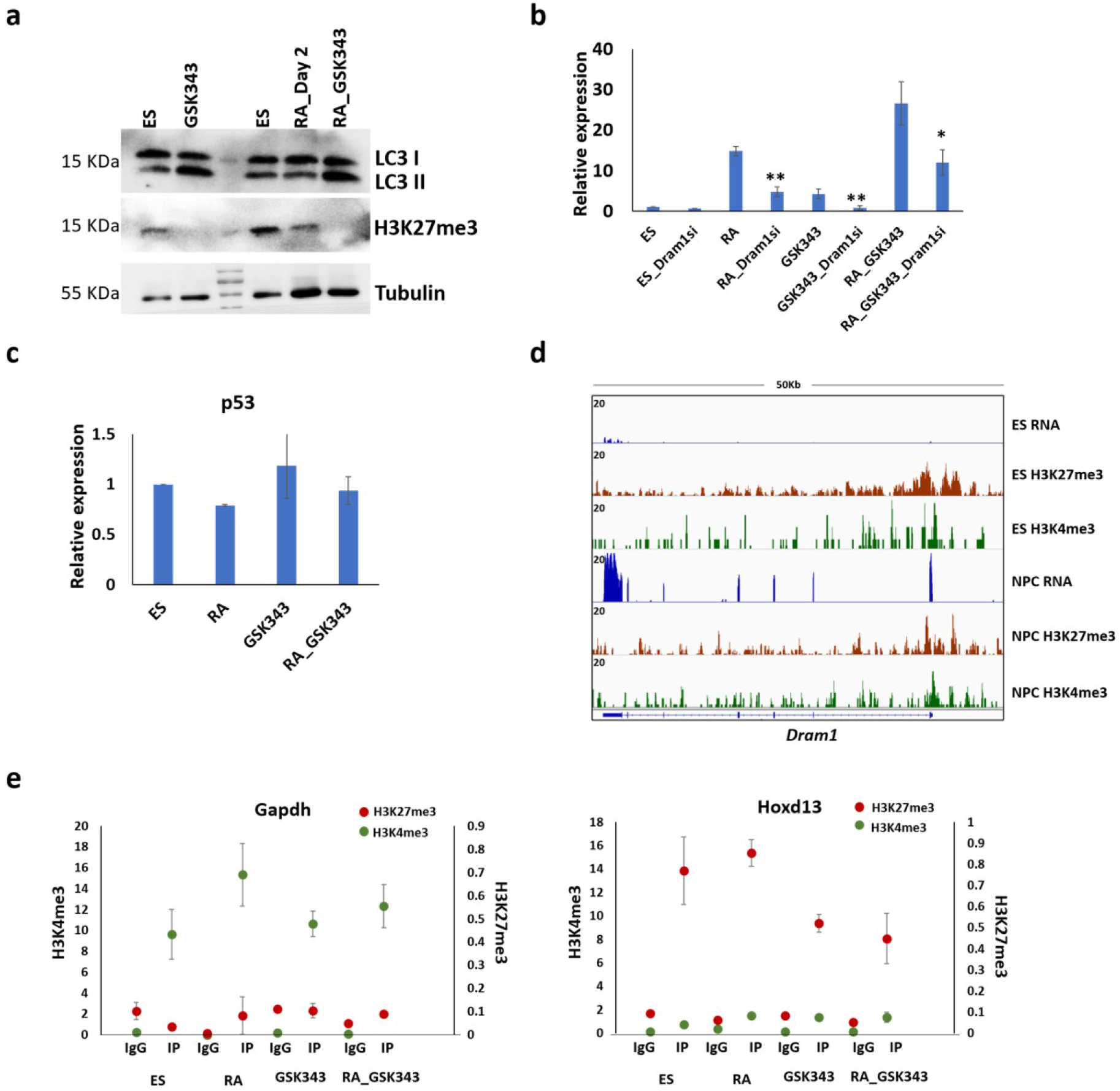
S3a: Immunoblot depicting LC3I, LC3II and H3K27me3 in mESCs, RA, GSK343 and RA_GSK343 treated cells. α tubulin is used as loading control. S3b: RT-qPCR analysis of *Dram1* expression in mESCs; RA, GSK343 and RA_GSK343 treated cells in presence of non-targeting siRNA and *Dram1* siRNA. Y axis represents fold change wrt *Gapdh* normalised to mESCs. N=3. Error bar indicates standard error of the mean (SEM). S3c: RT-qPCR analysis of *p53* expression in mESCs; RA, GSK343 and RA_GSK343 treated cells. Y axis represents fold change wrt *Gapdh* normalised to mESCs. N=3. S3d: IGV snapshot depicting RNA, H3K4me3 and H3K27me3 enrichment over the *Dram1* promoter in mESCs and NPCs. S3e: ChIP-qPCR analysis depicting the enrichment of H3K4me and H3K27me3 at the *Gapdh* and *Hoxd13* promoter. The Y axes represent percentage of input. N=3 biological replicates.

**Supplemental Table 1:** Drugs used for epigenetic inhibitor screen

| Target | Drug | Concentration (μM) | Treatment (Hours) |
| --- | --- | --- | --- |
| BRPF2(1)/TAF1(2) | BAY-299 | 1 | 1 |
| BAZ2A/B | BAZ2-ICR | 1 | 1 |
| BAZ2A/B | GSK2801 | 3 | 1 |
| CBP, EP300 bromo | ICBP112 | 3 | 1 |
| CBP, EP300 HAT | A-485 | 0.8 | 3 |
| CBP, EP300 HAT negative | A-486 | 0.8 | 3 |
| BET | JQ1 | 0.2 | 6 |
| BET (negative) | -JQ1 | 0.2 | 6 |
| BRD9 | I-BRD9 | 1 | 18 |
| PRMT type I | MS023 | 1 | 20 |
| SMYD2 | BAY598 | 1 | 24 |
| JMJD3, UTX, JARID1B | GSK J4 | 5 | 24 |
| WDR5 | OICR-9429 | 3 | 24 |
| WDR5 negative | OICR-0547 | 3 | 24 |
| SETD7 | R-PFI-2 | 1 | 24 |
| SETD7 (negative) | S-PFI-2 | 1 | 24 |
| SMYD2 | PFI-5 | 3 | 24 |
| PRMT3 | SGC 707 | 1 | 24 |
| PRMT3 | SGC XY | 1 | 24 |
| SUV420H1/2 | A-196 | 1 | 48 |
| LSD1 | GSKLSD1 | 1 | 48 |
| G9a, EHMT1 | A-366 | 1 | 72 |
| EED | A-395 | 1 | 72 |
| EED (Negative) | A-395N | 1 | 72 |
| EZH2 | GSK343 | 3 | 72 |
| Dot1L | SGC0946 | 1 | 72 |
| PRMT4 | TP-064 | 1 | 72 |
| PRMT4 (negative) | TP-064N | 1 | 72 |
| G9a, EHMT1 | UNC0642 | 1 | 72 |
| EZH2, EZH1 | UNC1999 | 3 | 72 |
| EZH2, EZH1 (negative) | UNC2000 | 3 | 72 |

**Supplemental Table 2:**
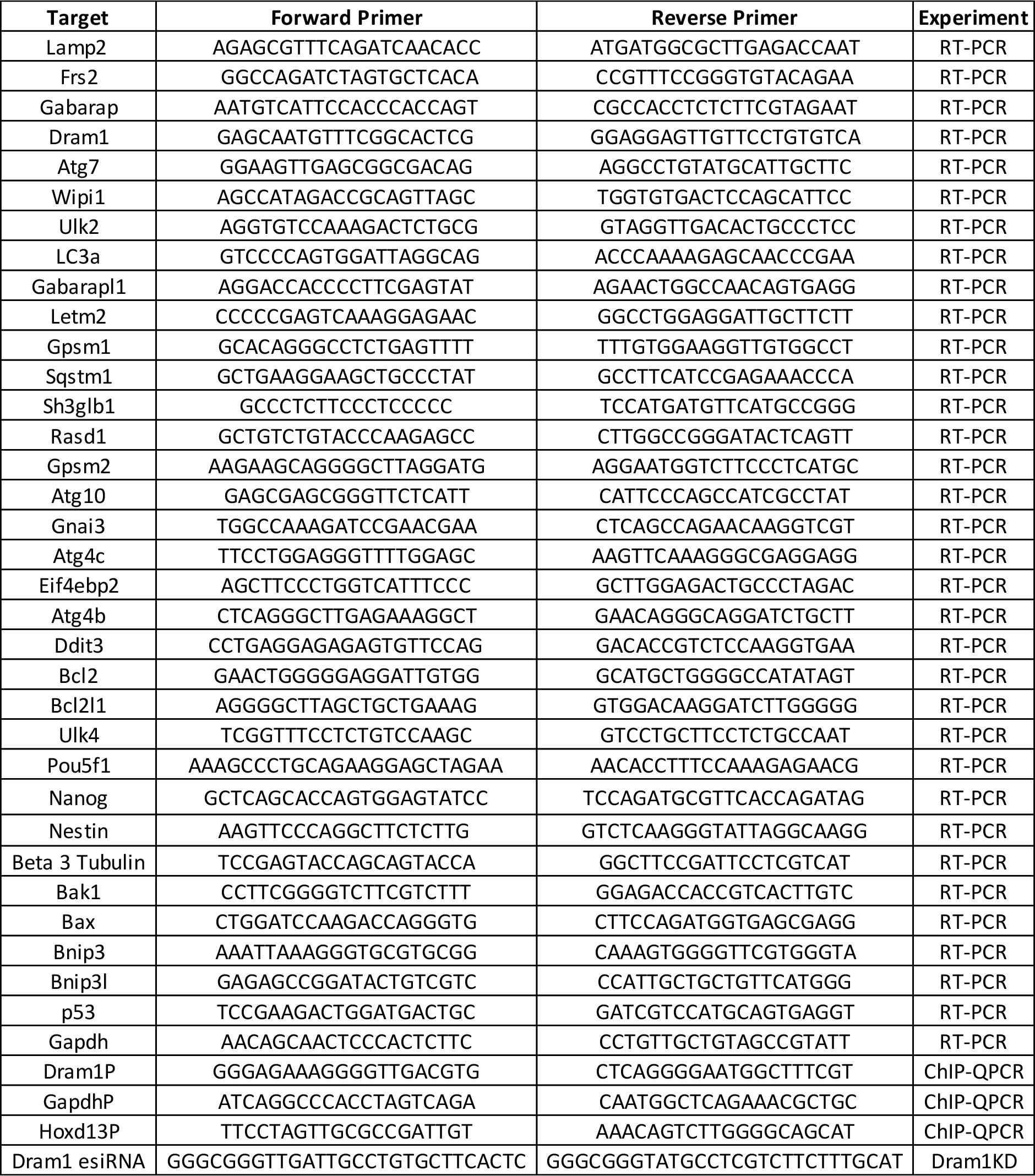
primers used for the study

